# Structural stability of SARS-CoV-2 degrades with temperature

**DOI:** 10.1101/2020.10.12.336818

**Authors:** A. Sharma, B. Preece, H Swann, X. Fan, R.J. McKenney, K.M. Ori-McKenney, S. Saffarian, M.D. Vershinin

## Abstract

SARS-CoV-2 is a novel coronavirus which has caused the COVID-19 pandemic. Other known coronaviruses show a strong pattern of seasonality, with the infection cases in humans being more prominent in winter. Although several plausible origins of such seasonal variability have been proposed, its mechanism is unclear. SARS-CoV-2 is transmitted via airborne droplets ejected from the upper respiratory tract of the infected individuals. It has been reported that SARS-CoV-2 can remain infectious for hours on surfaces. As such, the stability of viral particles both in liquid droplets as well as dried on surfaces is essential for infectivity. Here we have used atomic force microscopy to examine the structural stability of individual SARS-CoV-2 virus like particles at different temperatures. We demonstrate that even a mild temperature increase, commensurate with what is common for summer warming, leads to dramatic disruption of viral structural stability, especially when the heat is applied in the dry state. This is consistent with other existing non-mechanistic studies of viral infectivity, provides a single particle perspective on viral seasonality, and strengthens the case for a resurgence of COVID-19 in winter.

**Statement of Scientific Significance:** The economic and public health impact of the COVID-19 pandemic are very significant. However scientific information needed to underpin policy decisions are limited partly due to novelty of the SARS-CoV-2 pathogen. There is therefore an urgent need for mechanistic studies of both COVID-19 disease and the SARS-CoV-2 virus. We show that individual virus particles suffer structural destabilization at relatively mild but elevated temperatures. Our nanoscale results are consistent with recent observations at larger scales. Our work strengthens the case for COVID-19 resurgence in winter.

## Introduction

SARS-COV-2 is a virus of zoonotic origin which was first identified in humans in late 2019 [1]. Similar to other coronaviridae[2], the viral particles are enveloped and polymorphic decorated by a variable number of S protein spikes on their membrane[3]. One of the most confusing and yet urgently pressing questions at the time of this writing is whether the COVID-19 pandemic caused by SARS-CoV-2 will show seasonal character. Climate and seasonal dependence was expected early in the pandemic [4] due to similarity with other human coronavirus diseases [5], however the rates of infections have failed to strongly decline in the summer of 2020, leading to widespread doubts about COVID-19 seasonality. At the same time, a mounting body of evidence, from theoretical studies [6] to experimental research on viral populations and their infectivity [7,8] suggest that seasonality is indeed to be expected. However an understanding of how SARS-CoV-2 survives different environmental conditions is still incomplete and mechanisms of virus particle degradation are poorly mapped out. This then creates uncertainty for public health policy and its forward projection.

A key challenge in studying SARS-CoV-2 is the extreme level of threat associated with the live virus and the resultant need for high safety standards for such work. Aside from the envelope and S proteins, SARS-CoV-2 also packages the positive sense RNA genome encapsidated with thousands of copies of nucleocapsid, N proteins. SARS-CoV-2 also packages thousands of copies of matrix protein (M) which consists of three membrane spanning helixes with small intraluminal and extra luminal domains. In addition, an unknown number of envelope (E) proteins, which contain a single membrane spanning helix, are also packaged in each virion.

We have previously shown that similar to SARS-CoV [9], the expression of SARS-CoV-2 M, E, and S proteins in transfected human cells is sufficient for the formation and release of virus like particles (VLPs) through the same biological pathway as used by the fully infectious virus [10]. These VLPs faithfully mimic the external structure of the SARS-CoV-2 virus. The VLPs however, possess no genome and thus present no infectious threat which enables rapid studies with reduced safety requirements. The ability to produce non-infectious VLPs further enabled us to devise and rapidly validate novel strategies for manipulation of these particles, most notably via the addition of protein tags to the S and M proteins (these findings are detailed in a separate manuscript). Here, we report studies of VLPs subjected to variable temperature conditions before or after being immobilized and dried out on a functionalized glass surface. We show that exposure of VLPs to a mildly elevated temperature (34 C) for as little as 30 minutes is sufficient to induce structural degradation. The effect is weaker for particles exposed to elevated temperatures in solution and stronger for exposure in the dry state. Overall, these results provide insight into the viral seasonality of SARS-CoV-2.

## Results

During initial refinement of VLP purification strategies and associated VLP characterization [10], we have found that such particles remain stable for at least a week if stored in liquid buffer at near 0 °C conditions (on ice). We therefore examined whether they would remain stable at room temperature under dry conditions on a model surface (Fig. 1). Spytag-S VLPs were adhered to microtubules (MTs) via spycatcher-Kinesin and these complexes were abundantly retained on poly-L-lysinated glass surfaces. The shapes observed via AFM imaging at 22 °C (Fig. 1A) have nearly monodisperse sizes and only slight shape variability consistent with individual VLPs. Features indicative of envelope disruption and VLP aggregation are rare. Note that individual MTs are usually washed out during sample preparation leaving behind characteristic depressions in the poly-L-lysinated surface. In addition, surface clumping of poly-L-lysine is discernible in the AFM data as background height inhomogeneity. Identically prepared samples imaged via AFM at 34 °C under dry conditions (Fig. 1B) are harder to image stably due to high amount of background noise which obscures poly-L-lysine inhomogeneity and MT washout sites (likely due to loose debris on the surface – plausibly a by-product of particle degradations). Features consistent with intact VLPs are prominent relative to noise levels and hence easy to resolve (Fig. 2), but they are so exceedingly rare at 34 °C that they are considered outliers (Fig. 3). They were seen in large area surveys in which each particle is mechanically probed only a few times. Such particles do not survive intact during even a single detailed zoomed-in scan (Fig. 2). It takes 20-30 minutes to install the sample into the AFM, approach the surface and validate tip condition. Therefore mildly elevated temperature has a rapid effect on VLP integrity. VLPs incubated at 34 °C in solution and imaged at room temperature (Fig. 1C) survive better than particles at 34 °C under dry conditions but still often appear disrupted or aggregated in AFM imaging.

**Fig. 1.**
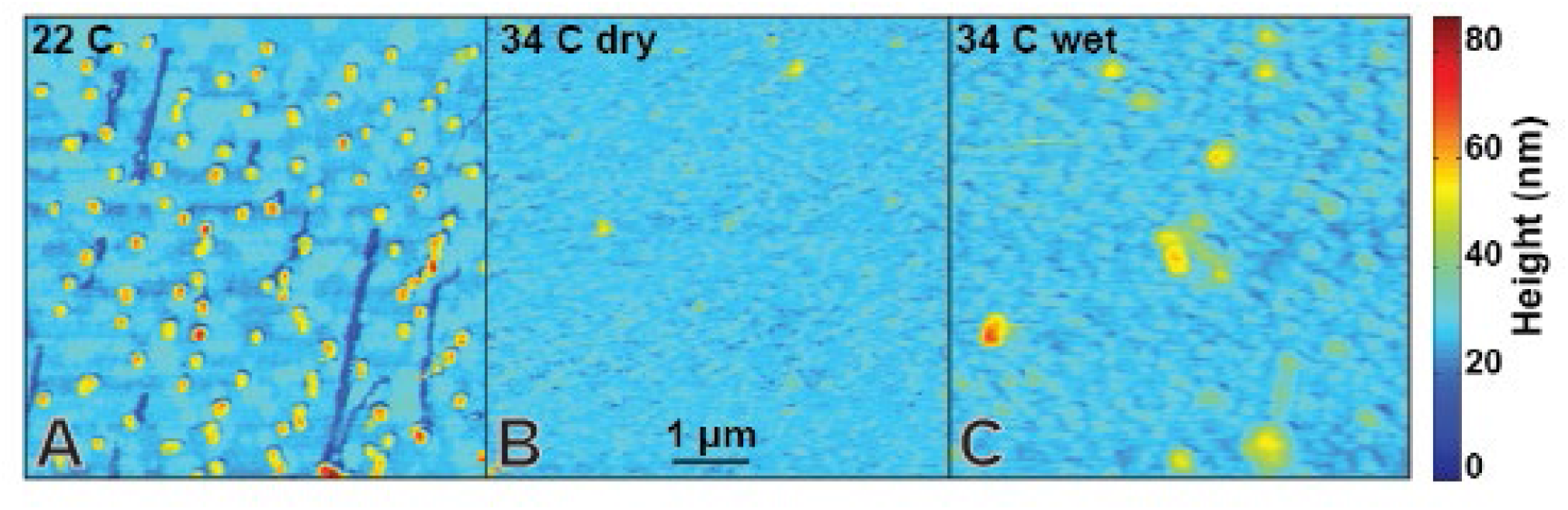
Sars-CoV-2 VLP stability as a function of environmental conditions. (A) VLPs are stable for hours on glass surfaces at room temperature under dry conditions. (B) VLPs imaged at 34 °C under dry conditions show high background noise and negligibly few features consistent with (A). MT washout sites can only be identified via high contrast enhancement (Fig. S1) and spatial peaks indicative of VLPs are rare and fragile (Fig. 2). (C) VLPs incubated at 34 °C in solution and imaged at room temperature are more consistent with (A) but also reveal widespread VLP disruption.

**Fig. 2.**
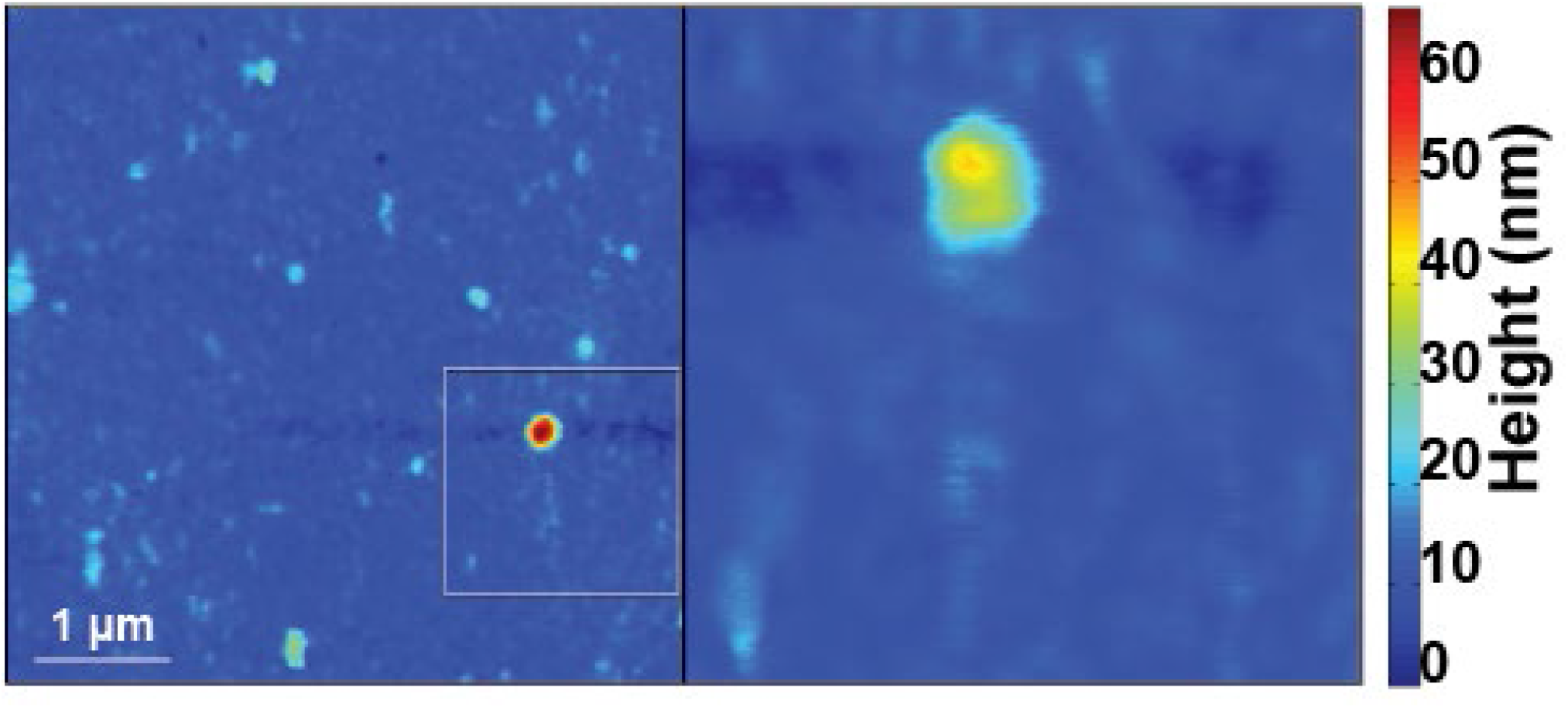
Sars-CoV-2 VLP disintegrates readily when AFM scanned at 34 C. Surface features consistent with individual VLPs are extremely rare. Even when such features are found in large area surveys (left), they cannot be scanned a second time, e.g. to obtain a zoomed in view (right). Zoom are is highlight on the left via a rectangle outline. The particles become flatter and show lateral spread consistent with mechanical VLP destruction [10].

**Fig 3.**
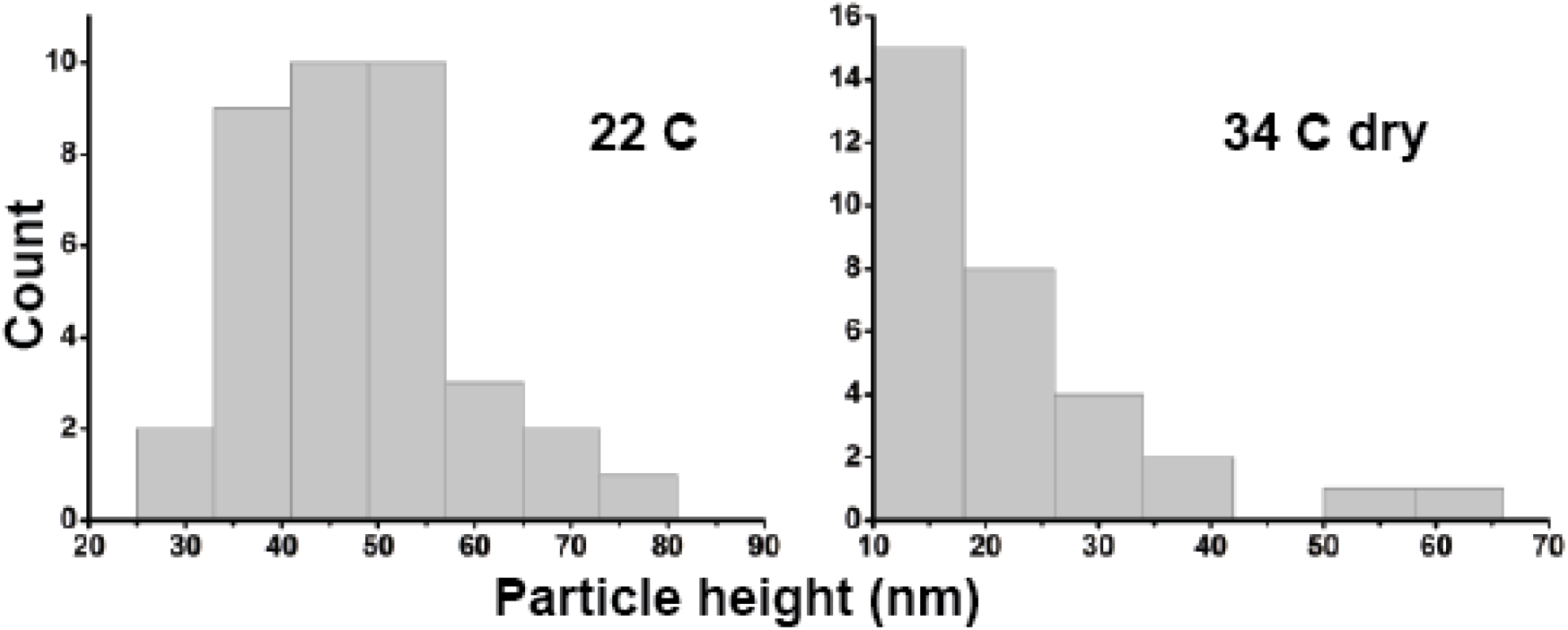
Temperature dependence of Z-height distributions of surface features. (A) Surface features have a strongly peaked height distribution consistent with air dried VLPs at 22oC. The median height of the particles is 43.4-51.4 nm (n=37). (B) Features observed at 34oC have median height 14.6-22.3 nm (n=31). CI estimated via bootstrap with 1e3 resamplings.

## Discussion

A common transmission route for SARS-CoV-2 is through bioaerosols created during sharp exhalation events such as sneezing or coughing. The bioaerosol droplets tend to dry out quickly due to high surface area and small volume [11,12]. Therefore virus particles may be exposed to both wet and dry conditions before coming into contact with and infecting the next host. It is widely recognized that virus particles often spread after their deposition on various surfaces [13] (although direct contact of the next host with contaminated bioaerosol may also be a viable route) and it is therefore also appreciated that virus particles can survive on various surfaces for an extended length of time[14].

The ability to make VLPs based on the SARS-CoV-2 genome, combined with abundant available structural information allowing for high precision design strategies for the VLPs, opens a unique opportunity for fast progress and allowed us to overcome the safety concerns associated with experiments on the full virus. Here we used this technology to study the stability of the viral envelope and associated proteins (M, E, and S) under different environmental conditions. As might be expected a priori, the VLPs do indeed degrade when exposed to elevated temperatures. Our AFM imaging revealed that negligibly few particles retain their shape and even those exceptional particles degraded nearly instantly during scanning and hence are likely already structurally impaired. The unexpected finding is how little heating it takes to degrade VLPs – just 34 °C was sufficient for a dramatic effect. Surfaces at 34 °C are warm to the touch but not hot. Such conditions are likely widespread in many locales for outdoors surfaces. In contrast, surfaces at 22 °C do not promote rapid VLP degradation, suggesting that surfaces commonly found indoors and the surfaces located outdoors during colder seasons may allow for prolonged viral survival and possibly extended viral spread. It is hard to estimate how all individual contributing factors would contribute to the epidemiological picture on the ground. Nonetheless, our findings draw parallels between the stability of SARS-CoV-2 and the original SARS viruses [15] and add to a growing body of research suggesting more viral spread is likely at lower temperatures via a variety of possible contributing factors [6].

## Materials And Methods

### Glass and MT preparation

Glass coverslips (VWR, Radnor, PA) were cleaned with dd-H_2_O and Ethanol. Clean slides were treated with 0.01% w/v Poly-L-Lysine (Millipore Sigma, Burlington, MA) for 30 min, washed and dried. Porcine microtubules (Cytoskeleton, Denver, Co) were polymerized according to manufacturer’s instructions with ~140mM of tubulin concentration. MTs were further diluted 30x in assay buffer (35mM PIPES, 5mM MgSo4, 1mM EGTA, 0.5mM EDTA, 10mM GTP and 40μm Taxol) at pH 7.2.

### VLP functionalization

VLPs suspended in PBS buffer were mixed at 4°C with KIF5B-spycatcher in (buffer) for 7-8 hours on a nutating mixer (VWR, Radnor, PA) to ensure Spy Tag-Spy Catcher binding. Different mixing concentrations of VLPs and KIF5B-spycatcher construct were tested. Excessively high concentration of KIF5B-spycatcher led to the formation of large aggregates of VLPs (~350-600nm lateral dimensions) while low concentrations of KIF5B-spycatcher led to reduced MT-VLP binding. Optimum binding of tagged S-protein with KIF5B-spycatcher resulted in predominantly single VLPs bound to MTs. MT were incubated with the KIF5B-spycatcher + VLP solution for 30 min at 4°C. VLPs attached on the MT were pelleted by spinning the solution at low speeds of 2500 rpm for 10 min in assay buffer. MT-VLP complexes were gently resuspended (buffer: 35mM PIPES, 5mM MgSo4, 1mM EGTA, 0.5mM EDTA, 10mM GTP and 40um Taxol) and then incubated on the Poly-L-Lysine functionalized glass surface for 20 min followed by H_2_O wash and gentle drying under nitrogen gas flow.

### AFM imaging and Temperature measurements

AFM experiments were performed using Dimension Icon AFM with an MLCT-BIO-DC probe (Bruker, Santa Barbara, CA, USA) in PeakForce QNM in air mode at room temperature (Scientific reports ref). The AFM was located on a dedicated optical table and featured a full acoustic enclosure. A small heater was placed in the enclosure to increase the temperature of the AFM chamber. Temperature was allowed to equilibrate prior to experiments and was measured using manufacturer calibrated NIST-traceable Thermo Hygrometer Barometer PCE-THB 40 (PCE Instruments, Jupiter, FL).

### Recombinant Protein purification

A constitutively active version of the human kinesin-1 motor truncation HsKIF5B(1-560) (K560 here after) was used in this work. To generate the K560-mScarlet-Strep-SpyCatcher construct, the coding sequences of human K560 (or its rigor mutant, K560-T92N), fluorophore mScarlet and Strep tag (WSHPQFEK) were amplified by PCR. The coding sequence of SpyCatcher was synthesized by IDT (Integrated DNA Technologies). These fragments were cloned into a pET28a-based vector using Gibson assembly. Flexible linkers (AAA, LE and GSGS) were introduced between these elements giving a final construct composed of K560-AAA-mScarlet-LE-StrepII-GSGS-SpyCatcher. All constructs were verified by DNA sequencing.

The recombinant proteins were expressed in BL21(DE3) cells in 2x YT medium. The cells were grown at 37 °C until an optical density at 600 nm (OD600) of 0.5, then induced with 0.2 mM isopropyl-β-D-thiogalactopyranoside (IPTG) 4 hours at 180C. Cells were resuspended in lysis buffer, containing protein buffer (50 mM Tris-HCl pH 8.0, 150 mM potassium acetate, 2 mM MgSO4, 1 mM EGTA, 5% glycerol) and freshly supplemented reagents (1 mM DTT, 1 mM ATP, 1 mM PMSF, protease inhibitor mix (Promega) and 1% Tween 20), then the cells were lysed using an Emusiflex C-3 (Avestin). To purify the proteins, cell lysates were treated with nuclease and incubated with Strep XT beads for 1 hour then washed with lysis buffer and eluted by elution buffer (protein buffer plus 100 mM biotin and 0.1% NP-40). The proteins were subsequently purified by anion exchange using a Mono Q column (GE Healthcare) in HB buffer (35 mM PIPES-KOH pH 7.2, 1 mM MgSO4, 0.2 mM EGTA, 0.1 mM EDTA) with salt gradient from 200 mM to 500 mM. To further purify the proteins, the Mono Q elutions containing desired proteins were further purified by concentration on Amicon Ultra filters followed by size-exclusion chromatograph using a Superdex 200 10/30 GL column (GE Healthcare Life Sciences) in GF150 buffer (25 mM HEPES pH 7.5, 150 mM potassium acetate, 1 mM MgCl2) with 0.01% NP-40.

## Acknowledgements

This work was supported by NSF RAPID 2026657 award to MV and SS, NIGMS grant 5R35GM124889 to RJM and NIGMS grant R35GM133688 to KMOM.

## Author Contributions

All authors designed research and co-wrote the manuscript; A.S., H.S., B.P. and performed research; X.F., K.M.O.-M. and R.J.M. contributed new reagents; M.V. and S.S. supervised research.

## Supplementary Information

**Fig. S1.**
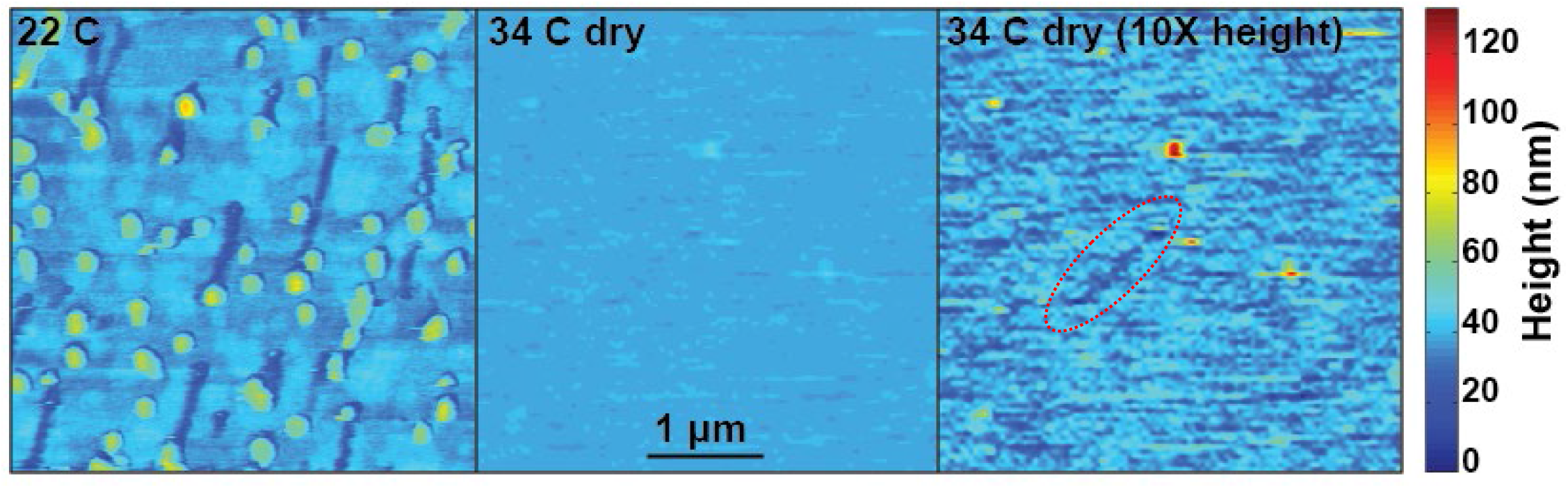
A comparison of features seen at 22 °C and 34 °C in 4 um x 4 um fields of view. VLPs and MT washout sites are readily identifiable for scans at room temperature, but are difficult to see with an identical colormap presentation due to elevated background noise. However, artificially extending the z range of the data helps reveal the presence of MT washout sites (right panel, washout site highlighted with red oval). Faint modulation in the rightmost image is most likely due to electronic noise in the imaging system although a mechanical vibration noise contribution cannot be excluded.

